# Inhibition of acid ceramidase as a therapeutic strategy for Niemann-Pick C disease

**DOI:** 10.1101/2024.11.13.623427

**Authors:** Isabelle Haynes, Simon E. Ward, D. Heulyn Jones, Emyr Lloyd-Evans, Helen Waller-Evans

## Abstract

Lysosomal storage disorders (LSDs) are a group of individually rare diseases that as a group constitute the most common forms of childhood neurodegeneration. These life limiting and life shortening diseases have the largest economic burden per patient of any rare disease with the majority being orphan diseases with no disease modifying therapy. Whilst there is significant development in the area of gene therapies for LSDs, these can only be used to treat individual diseases, are expensive, and in recent times many companies have abandoned their gene therapy pipelines. Small molecules on the other hand can be developed to target common mechanisms of pathogenesis or common central disease phenotypes providing therapies that can treat multiple diseases. In this study we report acid ceramidase as a novel therapeutic target for treating Niemann-Pick type C (NPC) disease. Using siRNA, and the literature tool compound carmofur, we observed a reduction in lysosomal area and reduced accumulation of the lipids cholesterol and LBPA in an NPC cellular model. These lipids are known to accumulate as secondary storage molecules across many lysosomal diseases, particularly primary sphingolipid storage disorders where lyso-sphingolipids are known to accumulate via the action of acid ceramidase. As acid ceramidase has also been shown to be a therapeutic target for Krabbe disease, our findings indicate that this is a novel target for the potential treatment of multiple LSDs with primary or secondary storage of sphingolipids.

## Introduction

Niemann-Pick C (NPC) disease is a rare autosomal recessive inherited neurodegenerative disease caused by mutations in one of either of two genes *NPC1* or *NPC2*^*1*^. These genes encode proteins of the same name that are involved in lysosomal cholesterol and lipid homeostasis^1^. NPC1 is a 13 transmembrane domain protein of the lysosomal limiting membrane whilst NPC2 is a soluble intralysosomal cholesterol shuttle^1^. Impairment in NPC1 or NPC2 function result in classical cellular hallmarks of intralysosomal storage of free unesterified cholesterol, sphingosine, sphingomyelin, lyso(bis)phosphatidic acid (LBPA) and glycosphingolipids (GSL) that occur in all NPC patient tissues^1^. Lipid storage is a direct consequence of loss of NPC1 or NPC2 function but is exacerbated by abnormalities in endocytosis, lipid trafficking and lysosomal Ca^2+^ signalling^1^.

Amongst the lysosomal storage disorders (LSDs), NPC is unique as it is the only one where the simple sphingoid base sphingosine has been shown to significantly accumulate^2^. Sphingosine is generated by desaturation of sphinganine during the formation of ceramide and is released within lysosomes by the action of acid ceramidase which catalyses the removal of the fatty acid^1^. Whilst the primary function of acid ceramidase is to cleave ceramide into sphingosine, the enzyme is also known to act on glycosphingolipids in other LSDs, presumably owing to their abundance within lysosomes of these cells, and to generate the so called lyso-glycosphingolipids (lyso-GSL), such as glucosylsphingosine and lyso-GB3 in Gaucher disease and Fabry disease respectively^3^. Sphingosine and lyso-GSLs are extremely potent and potentially toxic molecules that if not regulated correctly can have significantly detrimental properties to the cell. Sphingosine, and other sphingoid bases including lyso-GSLs, have been known to modulate protein kinase C signalling^4^, ion channel signalling^5^ and growth factor signalling^6^, they are implicated in numerous human pathologies including diabetes, neuropathy, cancer and neurodegenerative diseases^7,8^. Despite this significant involvement in human pathologies there are very few therapeutic molecules that exist that are able to regulate the levels or activity of acid ceramidase to alter sphingosine or lyso-GSL levels.

Genetic proof of concept that acid ceramidase is a viable therapeutic target has been demonstrated previously in Krabbe mice crossed with mice null for the acid ceramidase gene *ASAH1*^*9*^. Psychosine levels were returned to near normal with improvements seen in myelination, tremor, organ weight and motor function and an almost 50% increase in lifespan^9^. Further evidence of benefit was provided using the small molecule acid ceramidase inhibitor carmofur^9^. Together these data provide evidence to support the concept that the lyso-GSL ‘psychosine’ (galactosylsphingosine), rather than the parental molecule galactosylceramide, which accumulates due to the genetic defect in lysosomal galactosylceramidase activity in Krabbe disease, is the primary cause of pathogenesis, confirming the so-called ‘psychosine hypothesis’^9^. Whilst this has also been hypothesised for other LSDs, it is the understanding of these authors that only for Krabbe disease does evidence exist that inhibition of acid ceramidase activity leads to functional improvements^9^. Reduction in total sphingolipid content using the serine palmitoyltransferase inhibitor myriocin has been shown in the past to be beneficial in terms of restoring lysosomal Ca^2+^ levels in NPC disease human fibroblasts^2^. However, this cannot be solely attributed to a reduction in sphingosine as total GSL would also be reduced, which is the mechanism of action of one of the few approved therapies for NPC disease, miglustat^1^. We therefore chose to test the impact of acid ceramidase inhibition on NPC cellular models to confirm whether reduction in lysosomal sphingosine is indeed responsible for improvement in NPC disease phenotypes, and to gauge the potential utility of this novel therapeutic strategy in LSDs where sphingolipids are typically considered as being secondary storage material.

## Materials and Methods

### Cells

The glial cell line, prepared from the spontaneously occurring *Npc1*^*m1N*^ wild-type (*Npc1*^*+/+*^) and null (*Npc1*^*-/-*^) BALB/c mouse, has been described previously^2^. Cells were grown in culture as monolayers in Dulbecco’s modified Eagle’s medium (DMEM) supplemented with 10% foetal bovine serum (FBS) and 1% L-glutamine (complete medium) in a humidified 5% CO_2_ incubator at 37°C. During culture, spontaneous immortalisation occurred as an expected result of growth at high density^10^, these colonies were selected, maintained and characterised (manuscript in preparation) and are used in this report. Immortalised *Npc1*^*+/+*^ and *Npc1*^*-/-*^ glia retain all key NPC disease phenotypes.

### *ASAH1* silencing

Silencing RNA against *Asah1* (Silencer© siRNA 71420, ThermoFisher) was transfected into *Npc1*^*-/-*^ glia using JetPEI transfection reagent (Polyplus) at 5 nM and 50 nM using manufacturer’s protocol. Briefly, siRNA, resuspended in RNAse-free water, and JetPEI were diluted to the appropriate concentration in 150mM NaCl, mixed, allowed to incubate for 30 min at room temperature, and added dropwise to cells growing on glass coverslips in a 24 well plate. Negative control siRNA that does not target any known coding sequence in mouse (Mission, Sigma) was included (50nM only) to control for non-specific effects. After 72 hours, cells were fixed in 4% paraformaldehyde and stained as indicated.

### Fluorescence microscopy

Cells were seeded on Ibidi chamberslides and left overnight to adhere prior to treatment with the indicated concentrations of carmofur for the indicated times. Following drug treatments, cells were fixed with 4% paraformaldehyde for 10mins followed by three washes in Dulbecco’s Phosphate Buffered Saline (DPBS). Cells were stained with anti-LAMP1 (clone H4A3, ab25630, Abcam) antibody diluted 1:500, or anti-LBPA (Z-LBPA, Echelon) antibody diluted 1:500, both in blocking solution (1% bovine serum albumin (BSA) with 0.1% saponin in DPBS) and left for 24h at 4°C prior to 3 × 5 minute washes in DPBS. Cells were then incubated with DyLight488 conjugated goat anti-mouse IgG (Thermofisher) diluted 1:500 in blocking solution for 30 minutes at room temperature followed by 3 × 5min washes in DPBS. For staining of cholesterol, cells were incubated with filipin complex (187.5 μg/mL) in complete medium for 30 minutes at room temperature and washed twice in DPBS. All cells were subsequently imaged using a Leitz Dialux 22 microscope fitted with cooLED illumination, ORCA ER CCD fluorescence camera and micro manager 1.4 imaging software^11^. Area of staining per cell was quantified using ImageJ^12^.

## Results

To determine if a reduction in acid ceramidase activity could result in a normalisation of cellular phenotypes in the mutant Npc1 glia we treated the cells with an increasing concentration of carmofur for 48h. As can be seen, *Npc1*^*-/-*^ null glia present with classic hallmarks of NPC disease, namely an expansion of the lysosomal system, as illustrated by an increase in LAMP1 positive puncta, the accumulation of LBPA within lysosomes and the redistribution of cholesterol from the plasma membrane to punctate storage lysosomes (Fig. 1). Following treatment of *Npc1*^*-/-*^ glia with carmofur there is a clear and significant reduction at 30nM in lysosomal expansion (33% reduction) and lipid storage (LBPA and filipin back to wild-type levels). However, at 300nM only filipin is reduced and at 3μM, whilst there is no difference in any of the lipid phenotypes compared to the untreated control, we did observe a statistically significant increase in lysosomal expansion with LAMP1.

**Fig 1.**
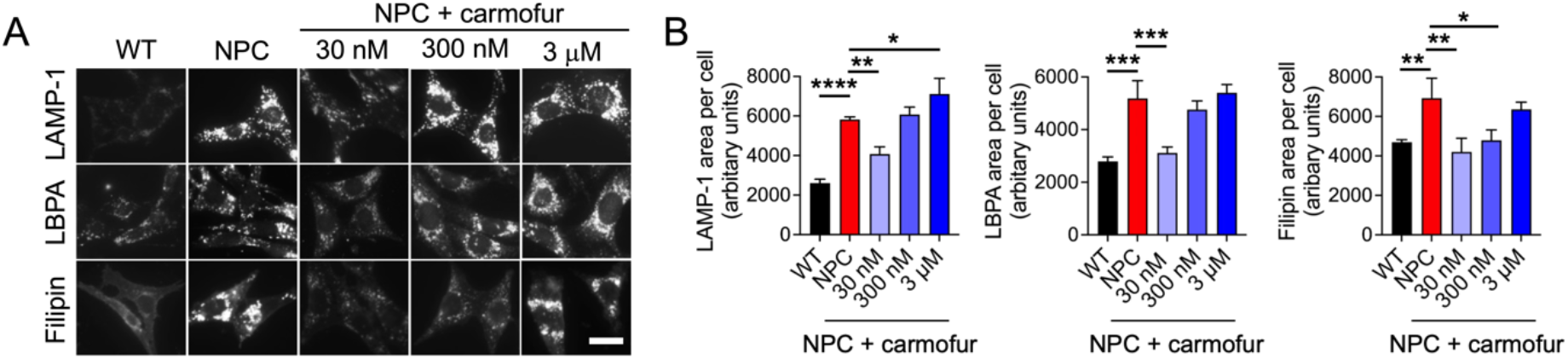
Effect of acid ceramidase inhibition with carmofur in *Npc1*^-/-^ null glia following a 48h incubation. *Npc1*^*-/-*^ glia (NPC) were treated for 48h with the indicated concentrations of carmofur and stained with filipin, to label cholesterol, and antibodies against LAMP-1 and LBPA, and compared to *Npc1*^*+/+*^ (WT) glia. Representative pictures shown, scale = 10μm (A). Graphs show quantification of staining area, n=3 (B). Data were analysed by one-way ANOVA followed by Tukey post-hoc testing, *p<0.05, **p<0.01, ***p<0.001, ****p<0.0001.

To confirm that the beneficial effects at the lower concentration of carmofur are due to reduction in acid ceramidase activity, rather than an off-target effect of carmofur, we treated *Npc1*^*-/-*^ glia with *Asah1* siRNA for 72h to reduce acid ceramidase levels and activity. As can be seen (Fig. 2), whilst the negative control and lower siRNA concentration of 5nM have almost no effect on lysosomal expansion or lipid accumulation, the higher concentration of 50nM *Asah1* siRNA did lead to reductions in all phenotypes. In comparison to the negative control siRNA, *Npc1*^*-/-*^ cells treated with 50nM *Asah1* siRNA had reduced lysosomal expansion (LAMP1 staining) by 33% and reduced lysosomal accumulation of LBPA by 25%. In comparison with the untreated *Npc1*^*-/-*^ glia, the 50nM *Asah1* siRNA treated cells did also demonstrate reduced lysosomal cholesterol accumulation (∼30% reduction in filipin staining). However, a reduction in filipin staining was also observed with the negative control siRNA (∼25%) but, significantly, not with the 5nM *Asah1* siRNA (9% reduction) suggesting a potential issue with the negative control.

**Fig 2.**
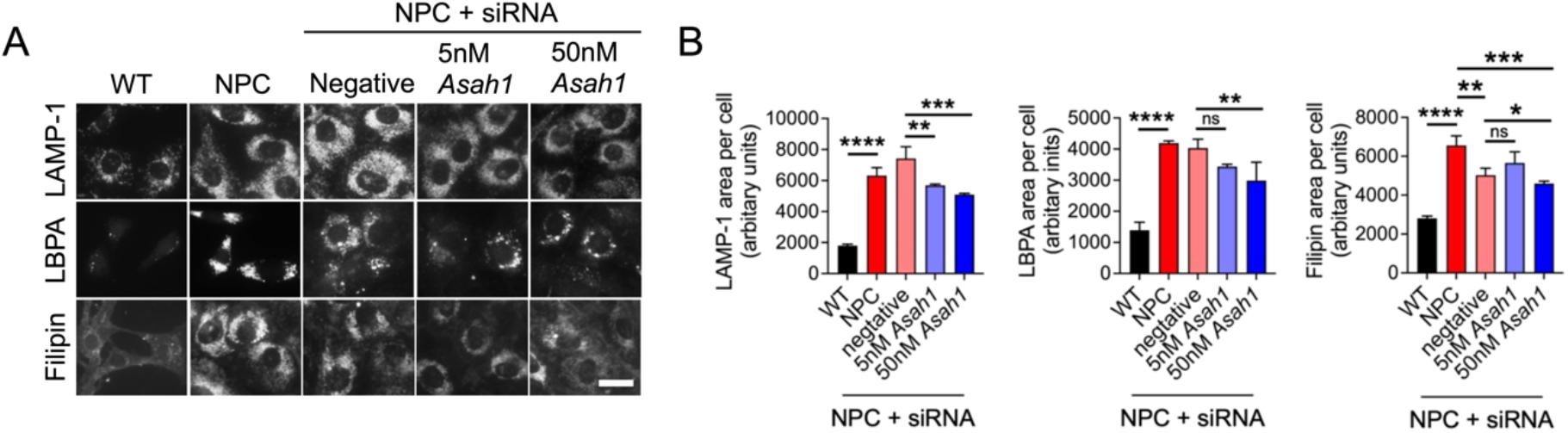
Knock down of acid ceramidase with *Asah1* siRNA in *Npc1*^*-/-*^ null glia provides partial correction of NPC phenotypes. *Npc1*^*-/-*^ glia (NPC) were treated for 72h with negative control siRNA (50nM) or siRNA targeting *Asah1* at the indicated concentrations and stained with filipin, to label cholesterol, and antibodies against LAMP-1 and LBPA, and compared to *Npc1*^*+/+*^ (WT) glia. Representative pictures shown, scale = 10μm (A). Graphs show quantification of staining area, n=3 (B). Data were analysed by one-way ANOVA followed by Tukey post-hoc testing, *p<0.05, **p<0.01, ***p<0.001, ****p<0.0001.

## Discussion

In this report we demonstrate for the first time that inhibition of acid ceramidase is a potential therapeutic strategy for NPC disease. Our findings provide both genetic and small molecule proof of concept that reducing acid ceramidase expression or inhibiting function can improve multiple phenotypes associated with the disease. Encouragingly, phenotypes including cholesterol storage, secondary lipid storage and lysosomal expansion were improved.

Furthermore, our data suggests that the beneficial effects observed previously with a 48h treatment of the serine palmitoyltransferase inhibitor myriocin in NPC patient fibroblasts^2^ are most likely due to a reduction in sphingosine rather than GSLs as the levels of these complex lipids are not modulated by carmofur^13^. Miglustat is the current standard of care for NPC and it is well documented that it takes 3-5 days for any beneficial effect of this drug to be seen in NPC cellular models, indicative of the longer turnover times for GSL storage within lysosomes^14^. Inhibition of acid ceramidase therefore provides a more rapid *in vitro* rescue of NPC lysosomal storage than miglustat, corresponding with the important role of sphingosine in the pathogenesis of NPC disease^2^ as, in comparison to carmofur, miglustat is unlikely to have any effect on lysosomal sphingosine storage.

Miglustat is a small molecule substrate reduction therapy that has been used in most territories worldwide as a disease modifying therapy for NPC disease for >15 years and most recently it has been approved for use in combination with arimoclomol in the US. Despite a clear benefit of miglustat on most NPC patients, leading to significant delay in disease onset, improved function and longer life expectancy^15^, it is well known that miglustat has no impact on lysosomal cholesterol levels in NPC cellular models or mouse models^14^. What impact this may have on the overall efficacy of miglustat in terms of improving NPC disease outcomes is unknown, as no therapy that has been shown to reduce lysosomal cholesterol in NPC disease models has yet translated into clinical efficacy. Our data indicate that inhibition of acid ceramidase, or reduction in expression, can reduce cholesterol accumulation in NPC cells, suggesting that this may be an improved therapeutic strategy in comparison to inhibition of glucosylceramide synthase and could lead to further therapeutic benefit for patients, particularly in combination with miglustat.

Finally, the presence of lyso-GSL storage across the majority of primary sphingolipidoses suggests that acid ceramidase could be a target for treating multiple LSDs. In addition to our data here, carmofur has been shown to be beneficial for treating Krabbe disease where it reduces storage of psychosine in the mouse model^9^. However, carmofur has numerous off target effects, it is a chemotherapy used in Japan to treat bowel cancer^16^ and is not an appropriate molecule to treat LSDs. Therefore, the development of a potent, titratable and selective acid ceramidase inhibitor, which is under development^17,18^, could be a novel pan-LSD therapeutic.

## Acknowledgments

The study was supported by Springboard award funding (SBF005/1129) from the Academy of Medical Sciences to HWE, Action Medical Research (GN3018) awarded to HWE, a Harrington Rare Disease Scholarship (GA_FFC03) awarded to HWE and an MRC IAA (524181/MB09) awarded to ELE.

## Author contributions

All experiments were performed by IH, experiments were designed by HWE, DHJ and ELE, manuscript was prepared by ELE, HWE, SW and DHJ.

